# Accurate and fast cell marker gene identification with COSG

**DOI:** 10.1101/2021.06.15.448484

**Authors:** Min Dai, Xiaobing Pei, Xiu-Jie Wang

## Abstract

Accurate cell classification is the groundwork for downstream analysis of single-cell sequencing data, yet how to identify marker genes to distinguish different cell types still remains as a big challenge. We developed COSG as a cosine similarity-based method for more accurate and scalable marker gene identification. COSG is applicable to single-cell RNA sequencing data, single-cell ATAC sequencing data and spatially resolved transcriptome data. COSG is fast and scalable for ultra-large datasets of million-scale cells. Application on both simulated and real experimental datasets demonstrates the superior performance of COSG in terms of both accuracy and efficiency as compared with other available methods. Marker genes or genomic regions identified by COSG are more indicative and with greater cell-type specificity.

## Introduction

With the broad application of various single-cell sequencing technologies, such as single-cell RNA sequencing (scRNA-seq)^1–3^ and single-cell assay for transposase-accessible chromatin using sequencing (scATAC-seq)^4–6^, as well as the rapid development of spatially resolved transcriptomics (spatial transcriptomics) technology^7–9^, how to accurately distinguish cells of interest from others or to characterize novel cell populations is becoming increasingly important^2,10,11^. The commonly used methods for cell marker gene identification usually rely on statistical tests to search for genes that are differentially expressed between cells of interest and all other cells in a dataset^12,13^. However, as statistical tests tend to identify candidates with systematic differences between two groups, when comparing one type of cells (target cells) with multiple other types of cells (non-target cells), the top-ranked differentially expressed genes selected by statistical methods may not be real cell markers. For example, a gene could be highly expressed in target cells and a small group of non-target cells, but almost non-detectable in other cells. Such gene could be selected as a marker gene for the target cells by expression-based statistical methods, but it could bring false results when being used for cell type characterization. Problematically, expression-based statistical methods are the default approaches for marker gene identification in most single-cell data analysis toolkits, including the two most commonly-used software, namely Scanpy^14^ and Seurat^15^.

Cosine similarity measures the relationship of two *n*-dimensional vectors using the cosine value of the angle between the vectors in the vector space. Unlike Euclidean distance which measures the positional difference between two vectors, cosine similarity compares the orientations of two vectors, which means if two genes have identical expression patterns but different expression abundance among a group of cells, the two genes will be considered as equivalent. Therefore, cosine similarity is scale-independent^16^ and is more sensitive to identify genes specifically expressed in target cells, yet it has not been applied on cell marker gene identification so far.

As the single-cell RNA-seq technology becomes more mature and popular, the number of cells captured by each experiment is rapidly increasing^1^, yet the currently available cell marker gene identification methods often suffer from their slow speed when handling data with a large number of cells. In addition, with the development of scATAC-seq^4–6^ and spatial transcriptomics technologies^7–9^, the need for a universal method with the capability to identify cell marker genes from multiple types of single-cell data modalities is rapidly emerging.

To address the challenges mentioned above, we developed COSG (COSine similarity-based marker Gene identification), a method to identify cell marker genes with better accuracy and faster speed. COSG outperforms existing tools in terms of the expression specificity of identified marker genes and the analysis time needed for large-scale datasets. In addition to scRNA-seq data, COSG can also be applied to scATAC-seq and spatial transcriptome data with good performance. Therefore, COSG can serve as a general method for cell marker gene identification across different data modalities to facilitate downstream analysis and discoveries.

## Results

### COSG uses cosine similarity to evaluate the expression specificity of genes

The basic concept of COSG is to compare the expression patterns of two genes within a given cell population by evaluating the angles between the vectors representing the expression of each gene in an *n*-dimensional cell space. Within the cell space, each dimension represents a cell. The representing vector for each gene consists of *n*-basis (*n* equals to the number of total detected cells), and the coordinate of each basis represents the gene’s expression level in each cell. Therefore, the cosine similarity of two genes equals the cosine value of the angle between each gene’s representative vector in the cell space. The more similar the expression patterns, the smaller the angle is. If two genes have identical expression patterns, the angle between their representative vectors will be zero, regardless of their abundance difference.

The marker gene identification process of COSG starts with multiple groups of cells pre-classified by other single-cell analysis tools. To identify marker genes for each cell group, COSG first creates an artificial gene (***λ***_***k***_) which only expresses in cells of a given group, e.g., Group *k* (*G*_*k*_, *k* ∈ {1, …, *K*}) and does not express in any other groups of cells, thus ***λ***_***k***_ would be the ideal marker gene for cells belonging to *G*_*k*_ (Fig. 1). The representative vector for each expressed gene (***g***_***i***_, *i* ∈ {1, …, *M*}) will be compared with the representative vector of ***λ***_***k***_, genes whose representative vectors with the smallest angles to the representative vector of ***λ***_***k***_ and the largest angles to the representative vectors of other cell groups (***λ***_***t***_, *t* ∈ {1, …, *K*} and *t* ≠ *k*) will be selected as the marker genes for *G*_*k*_. Here, we define COSG score as 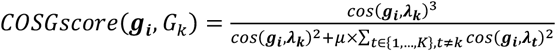, where *cos*() calculates the cosine similarity between the representative vectors of two genes, and *μ* (*μ* ≥0) is a user-defined hyperparameter as the penalty score (by default, *μ* = 1). The output of COSG is a list of candidate marker genes starting with the ones with the highest COSG scores for each cell group. COSG is available both in Python and R, and can be seamlessly used with Scanpy^14^ and Seurat^15^.

**Fig. 1.**
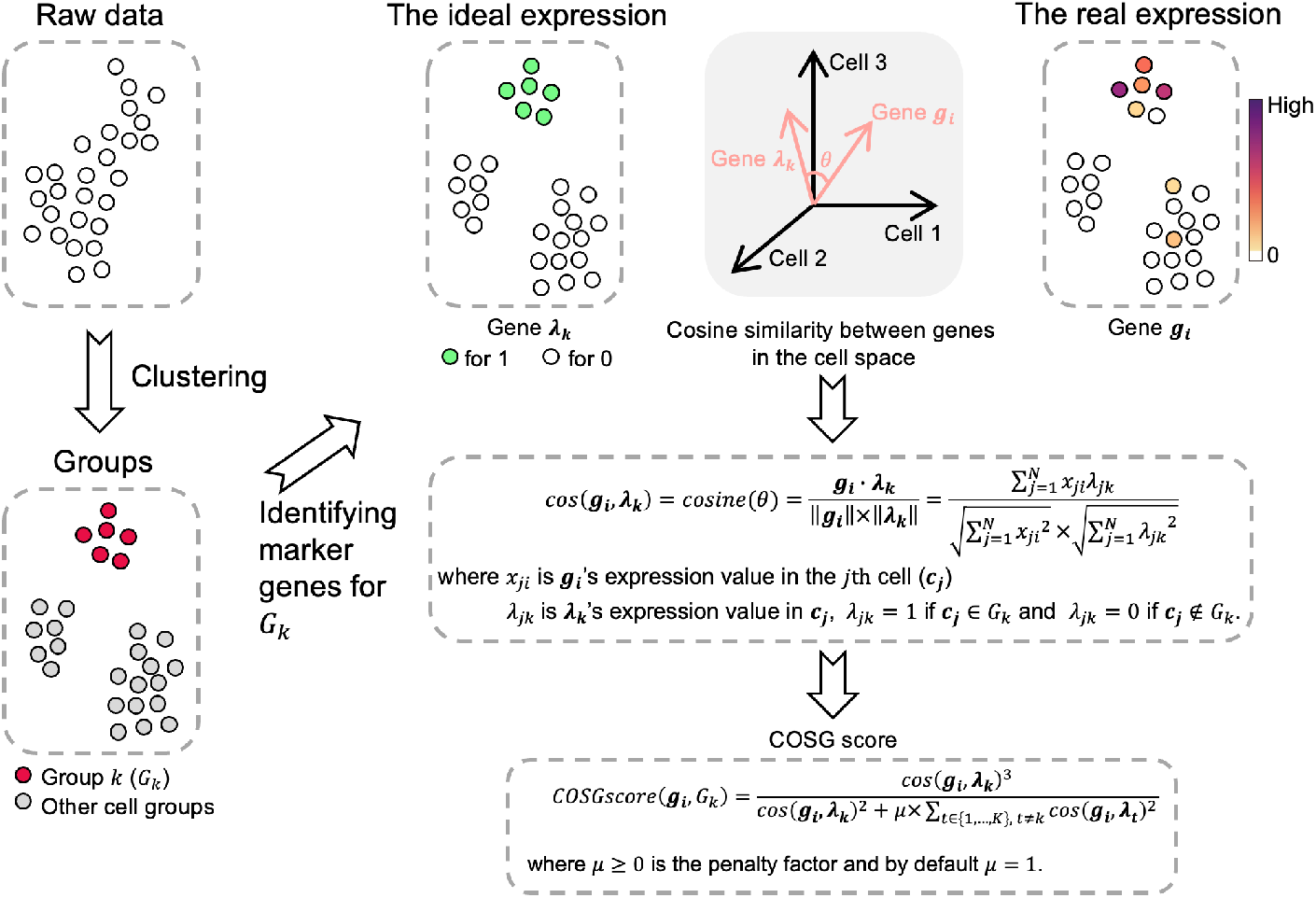
Workflow of COSG. The basic idea of COSG is to identify marker genes within a given group of cells by comparing the cosine values of the angles between the representative vectors of each detected gene and the assumed ideal marker gene. The input data of COSG should be normalized and clustered scRNA-seq data/scATAC-seq data/spatial transcriptome data. For a dataset of *N* cells (clustered into *K* groups) with *M* expressed genes, to identify marker genes for group *k* (*G*_*k*_, *k* ∈ {1, …, *K*}), COSG first creates an ideal marker gene ***λ***_***k***_ for *G*_*k*_, which was only detected in cells of *G*_*k*_ with uniformed expression value but not in any other group of cells. To examine whether a detected gene, ***g***_***i***_, *i* ∈ {1, …, *M*}, is a good marker gene for *G*_*k*_, COSG evaluates the expression similarity between gene ***g***_***i***_ and gene ***λ***_***k***_ among all cells by calculating the cosine values of the angles formed by the representative vectors of ***g***_***i***_ and ***λ***_***k***_ in the *N*-dimensional space spanned by all cells, then generates COSG score to reflect the expression specificity of ***g***_***i***_ in *G*_*k*_ by comparing the expression values of ***g***_***i***_ and ***λ***_***k***_ as well as ***λ***_***t***_ (*t* ∈ {1, …, *K*} *and t* ≠ *k*). As ***λ***_***t***_ represents the ideal marker genes for cell groups other than *G*_*k*_, the COSG score reflects the suitability of ***g***_***i***_ to serve as a marker gene for *G*_*k*_. By repeating the above procedures, COSG could identify marker genes for each group of cells.

### COSG identifies more indicative marker genes in scRNA-seq data

To test the function of COSG, we first generated 30 simulated scRNA-seq datasets with ground truth for known marker genes (Methods, Supplementary Table 1), and compared the performance of COSG with other 10 popular methods in the commonly used toolkits Scanpy^14^ and Seurat^15^ (Supplementary Table 2) on these datasets. We calculated the average overlapping ratios between the top 20 marker genes identified by each method and the true 20 marker genes of the 30 simulated datasets. The results showed that COSG outperformed all other tested methods (Fig. 2a and Supplementary Fig. 1).

**Fig. 2.**
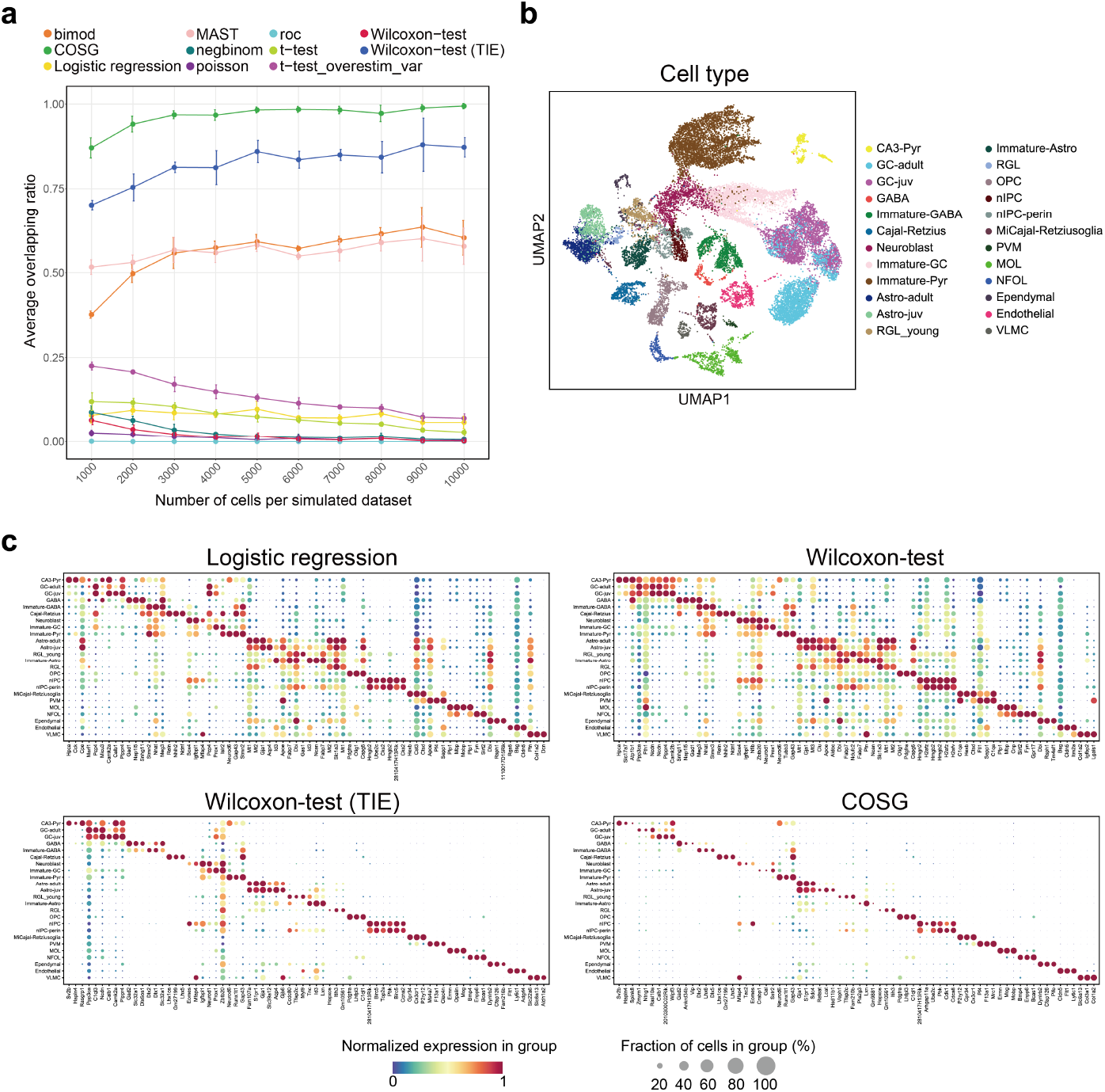
Performance comparison of COSG with other methods on scRNA-seq data. **a**, Average overlapping ratios of the top 20 marker genes identified by COSG or other 10 popular methods vs. the top 20 known marker genes of the 30 simulated datasets. Error bars represent the standard deviation of 3 datasets. **b**, UMAP projection of the scRNA-seq data of dentate gyrus cells from perinatal, juvenile, and adult mice. **c**, Expression dot plots of the top 3 marker genes identified by Logistic regression, Wilcoxon-test, Wilcoxon-test (TIE) and COSG for each cell type.

We then compared COSG with three well-used methods in Scanpy^14^, namely Logistic regression^17^, Wilcoxon Rank Sum test (Wilcoxon-test, also known as Mann-Whitney *U* test)^18,19^, and Wilcoxon-test with tie correction (denoted as Wilcoxon-test (TIE)) using two reported scRNA-seq datasets (Supplementary Table 3). Wilcoxon-test is the default method used in Seurat. It is also included in Scanpy and is the most widely used method for marker gene identification from scRNA-seq data. As scRNA-seq data usually contain many zero values (tied values), tie-correction is also implemented for Wilcoxon-test in Seurat. However, the default Wilcoxon-test in Scanpy does not perform tie correction and has been widely used by many published studies^20^ and cell atlas projects^21,22^.

The two reported scRNA-seq datasets used in this study were published by Hochgerner et al.^23^ and Stewart et al.^24^, respectively (Supplementary Table 3). The Hochgerner dataset contains 23,025 cells (belonging to 24 cell types) from the dentate gyrus tissue of perinatal, juvenile, and adult mice^23^. UMAP projection results confirmed the gene expression similarities among cells within each group (Fig. 2b). We examined the top 3 marker genes of each cell type identified by COSG and other methods, and found that most marker genes identified by Logistic regression or Wilcoxon-test are not cell type-specific (Fig. 2c). Wilcoxon-test (TIE) works slightly better, but still identified more non-specific marker genes as compared with COSG (Fig. 2c). About 54% marker genes (top 3 for each group) identified by COSG were also reported by Wilcoxon-test (TIE), but only 16% or 8% of them were identified by Logistic regression or Wilcoxon-test, respectively (Supplementary Fig. 2a). Expression pattern examination also revealed that, the top 3 marker genes for adult granule cells (GC-adult) identified by other methods almost all had relatively high expression abundance in at least one type of non-target cells, such as hippocampus CA3 pyramidal layer cells (CA3-Pyr) and juvenile GC cells (GC-juv), which are highly similar to GC-adult cells, yet 2 out of 3 marker genes identified by COSG had GC-adult cell-specific expression (Supplementary Fig. 2b). The Stewart scRNA-seq dataset contains 40,268 cells (belonging to 27 cell types) from human adult kidney tissue^24^, of which some cell types, especially those of the immune cells, were less distinguishable from each other by UMAP projection (Supplementary Fig. 3a). Again, the marker genes for almost all cell types identified by COSG showed high specificity, yet the other methods failed to reach the same standard (Supplementary Fig. 3b and 3c).

### COSG outperforms existing methods on large-scale datasets

To evaluate the computational performance and scalability of COSG, we measured the running time of COSG and the other 10 methods mentioned above (Supplementary Table 2) on 14 scRNA-seq datasets with cell numbers ranging from 1,000 to 150,000 (Supplementary Table 4). When handling scRNA-seq data of less than 10,000 cells, COSG and five other methods (namely t-test, t-test_overestim_var, Wilcoxon-test, Logistic regression, Wilcoxon-test (TIE)) finished the analysis almost instantly (Fig. 3a). Further comparison of these six methods on larger datasets with 10,000 to 150,000 cells demonstrated that COSG ran much faster than other methods, especially when the number of cells reached 150,000 (Fig. 3b and Supplementary Table 5). In addition, COSG identifies marker genes for over 1 million cells (1,331,984 cells) belonging to 37 cell types in less than 2 minutes (Supplementary Fig. 4).

**Fig. 3.**
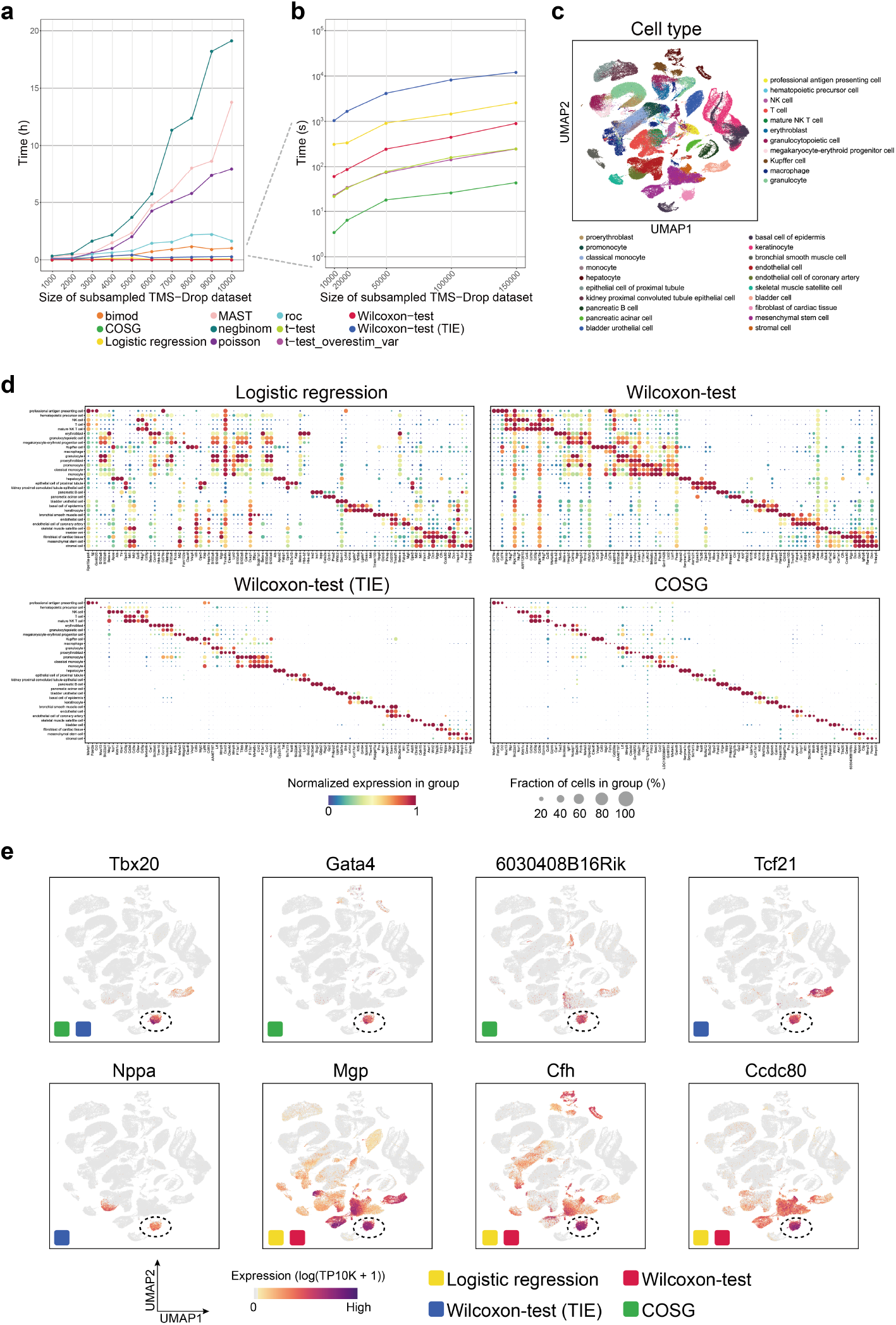
COSG efficiently and accurately identifies more indicative marker genes in large-scale scRNA-seq datasets. **a**, Running time of COSG and other 10 popular methods on subsampled Drop-seq datasets with cell numbers ranging from 1,000 to 10,000. **b**, Running time of the six fastest methods, namely COSG, t-test, t-test_overestim_var, Wilcoxon-test, Logistic regression and Wilcoxon-test (TIE) on subsampled Drop-seq datasets with cell numbers ranging from 10,000 to 150,000. **c**, UMAP projection of the scRNA-seq data with 150,000 subsampled cells. **d**, Expression dot plots of the top 3 marker genes identified by Logistic regression, Wilcoxon-test, Wilcoxon-test (TIE) and COSG for each group. **e**, Expression patterns of the top 3 marker genes for fibroblast of cardiac tissue identified by Logistic regression, Wilcoxon-test, Wilcoxon-test (TIE) and COSG. Cells classified as fibroblasts of cardiac tissue are indicated by dashed circles.

To examine whether the high efficiency of COSG is achieved without sacrificing its accuracy, we further analyzed the expression of the top 3 maker genes of each cell type identified by Logistic regression, Wilcoxon-test, Wilcoxon-test (TIE) and COSG from the above-mentioned 150,000 cells. Among the 31 cell types in this dataset, some cell types were difficult to be distinguished from each other by UMAP projections (Fig. 3c) or by marker genes identified by Logistic regression or Wilcoxon-test (Fig. 3d). Both COSG and Wilcoxon-test (TIE) reported specific marker genes for most cell types, but the processing time used by COSG was only 1/280 of that used by Wilcoxon-test (TIE), and the marker genes identified by COSG also had higher expression specificity (Fig. 3b and 3d, Supplementary Table 5). For example, the top 3 marker genes for fibroblast of cardiac tissue identified by COSG were all dominantly expressed in the target cells, whereas the top 3 marker genes identified by other methods all had high expression in one or more types of non-target cells (Fig. 3e). Taken together, these results demonstrated the advantages of COSG in handling large-scale datasets.

### COSG correctly identifies cell-type-specific marker regions in scATAC-seq data

We next assessed the performance of COSG on scATAC-seq data, which are much sparser and contain 10-20 times more features than scRNA-seq data^25^. Again, we compared the results generated by Logistic regression, Wilcoxon-test, Wilcoxon-test (TIE) and COSG using two reported scATAC-seq datasets (Supplementary Table 3). The first dataset (the Pijuan-Sala dataset) contains 301,316 detected genomic regions of 19,453 single nuclei from mouse embryos at 8.25 days post-fertilization^4^. The second dataset contains 451,999 detected genomic regions of 33,819 bone marrow and peripheral blood mononuclear cells (BMMCs and PBMCs, respectively) from healthy human donors^6^. We first examined the computational efficiency of COSG on scATAC-seq data. A broad cell type annotation (17 cell types) and a fine cell type annotation (23 cell types) were applied to the ATAC-Granja dataset. In all cases, COSG consumed less than 2 minutes, whereas Logistic regression and Wilcoxon-test were about 30 times slower than COSG, and Wilcoxon-test (TIE) was more than 300 times slower than COSG (Fig. 4a, Supplementary Table 6).

**Fig. 4.**
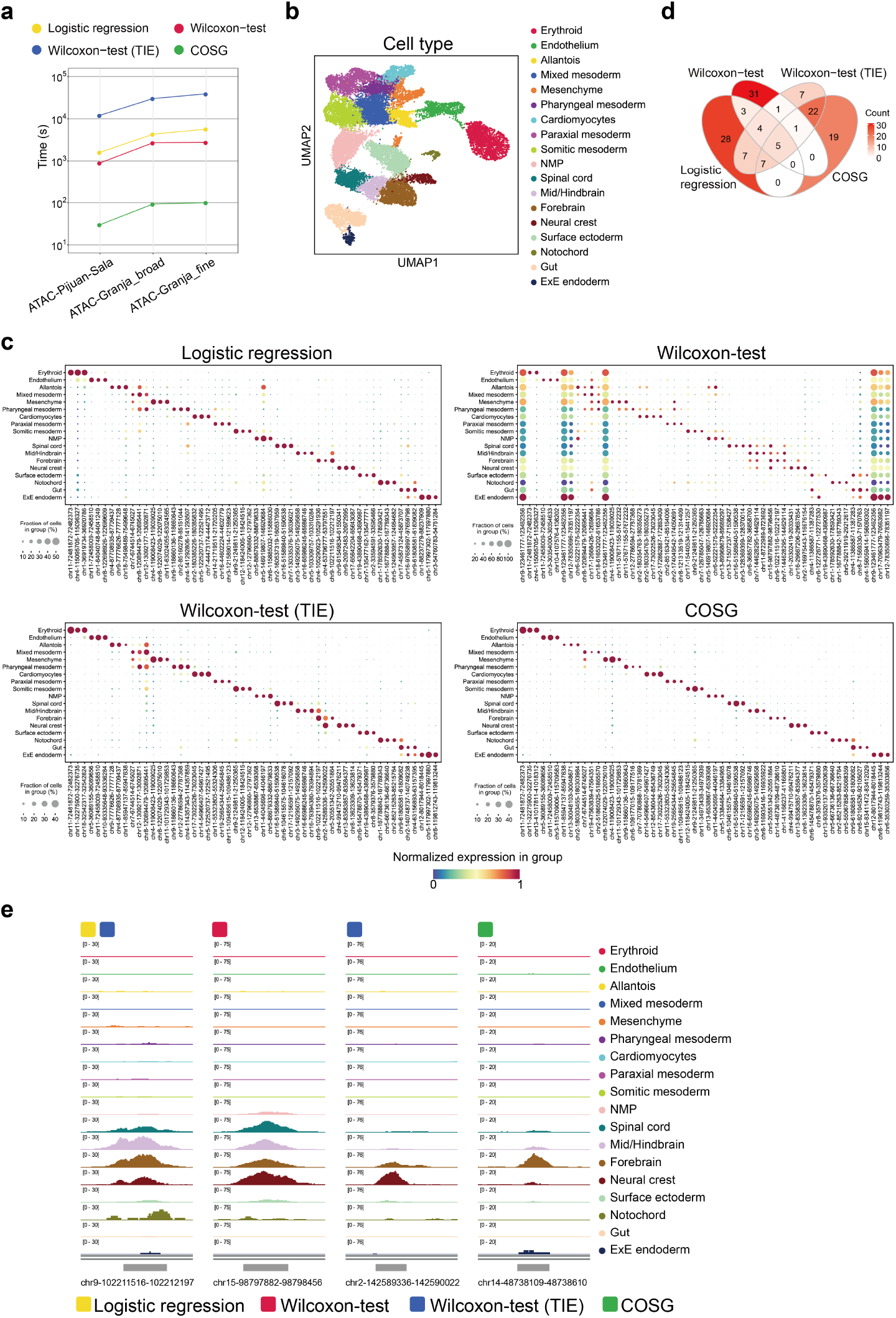
COSG outperforms existing methods on scATAC-seq data. **a**, Running time of Logistic regression, Wilcoxon-test, Wilcoxon-test (TIE) and COSG on three scATAC-seq datasets of different sizes and groups. **b**, UMAP projection of the scATAC-seq data from 19,453 mouse embryonic cells at 8.25 days post-fertilization (the ATAC-Pijuan-Sala dataset). **c**, Expression dot plots of the top 3 marker regions identified by Logistic regression, Wilcoxon-test, Wilcoxon-test (TIE) and COSG for each cell type. **d**, Venn diagram of the joint set of the top 3 marker regions for each cell type identified by different methods. **e**, Normalized genome browser tracks of representative marker regions for forebrain cells identified by different methods. Each track represents the aggregated signals for all cells of the corresponding cell type.

The UMAP projection result of the Pijuan-Sala dataset^4^ shows overlaps of some cell types, especially the ones from undifferentiated mesoderm (Fig. 4b), which made marker gene identification more difficult. Similar to the results of scRNA-seq data, the top 3 marker regions identified by COSG were more specific than the ones identified by other methods (Fig. 4c). Majority of marker regions reported by COSG were not identified by Logistic regression or Wilcoxon-test (Fig. 4d). Taking forebrain cells as an example, the genomic region ‘chr14-48738109-48738610’ had specific accessibility in forebrain cells and was identified as one of the top 3 marker regions only by COSG, yet the marker regions identified by other methods showed high accessibility in non-forebrain cells, namely spinal cord, mid/hindbrain cells, or neural crest cells (Fig. 4e). Notably, region ‘chr2-142589336-142590022’, one of the top 3 marker regions for forebrain cells identified by Wilcoxon-test (TIE), showed much higher accessibility in neural crest cells than in forebrain cells (Fig. 4e).

Immune cells, especially subtypes of the same immune cell type (e.g., naive CD4^+^ T cell and memory CD4^+^ T cell), are usually highly similar to each other in terms of molecular features. Analysis results of both the broad cell-type annotation (including 17 cell types) and fine cell-type annotation (including 23 cell types) of the Granja scATAC-seq dataset showed that, marker regions identified by COSG had higher cell type specificity than the ones identified by other methods, especially for different T cell subtypes (Supplementary Fig. 5 and Supplementary Fig. 6).

### COSG holds advantage in analyzing spatial transcriptome data

Spatial transcriptome data has emerged as a new data type in recent years, and the analysis of spatial transcriptome data also relies on marker gene identification to characterize cell types. To test the applicability of COSG on spatial transcriptome data, we first applied it on a dataset (the Spatial-brain_sagitta dataset, Supplementary Table 3) generated by 10x Genomics Visium platform using adult mouse brain. A total of 3,355 signal spots were detected in this dataset and clustered into 11 groups according to their gene expression profiles (Fig. 5a and 5b). To examine the accuracy of COSG, we compared the top 3 marker genes of each group identified by different methods (Fig. 5c). It is apparent that most marker genes identified by Logistic regression or Wilcoxon-test do not have cell type specificity. Wilcoxon-test (TIE) works better, but still picked up more non-specific cell markers as compared with COSG (Fig. 5c). We further examined the spatial expression pattern of Cluster 0’s top 3 marker genes identified by each method (Fig. 5d). The results showed that marker genes identified by COSG had higher and more specific expression among Cluster 0 cells as compared to markers identified by other methods. Similarly, application of the above-mentioned four methods on another 10x Genomics Visium dataset (the Spatial_brain_coronal dataset, Supplementary Table 3) generated using the coronal region of a mouse brain also demonstrates the capability of COSG in identifying more indicative marker genes from noisy data (Supplementary Fig. 7).

**Fig. 5.**
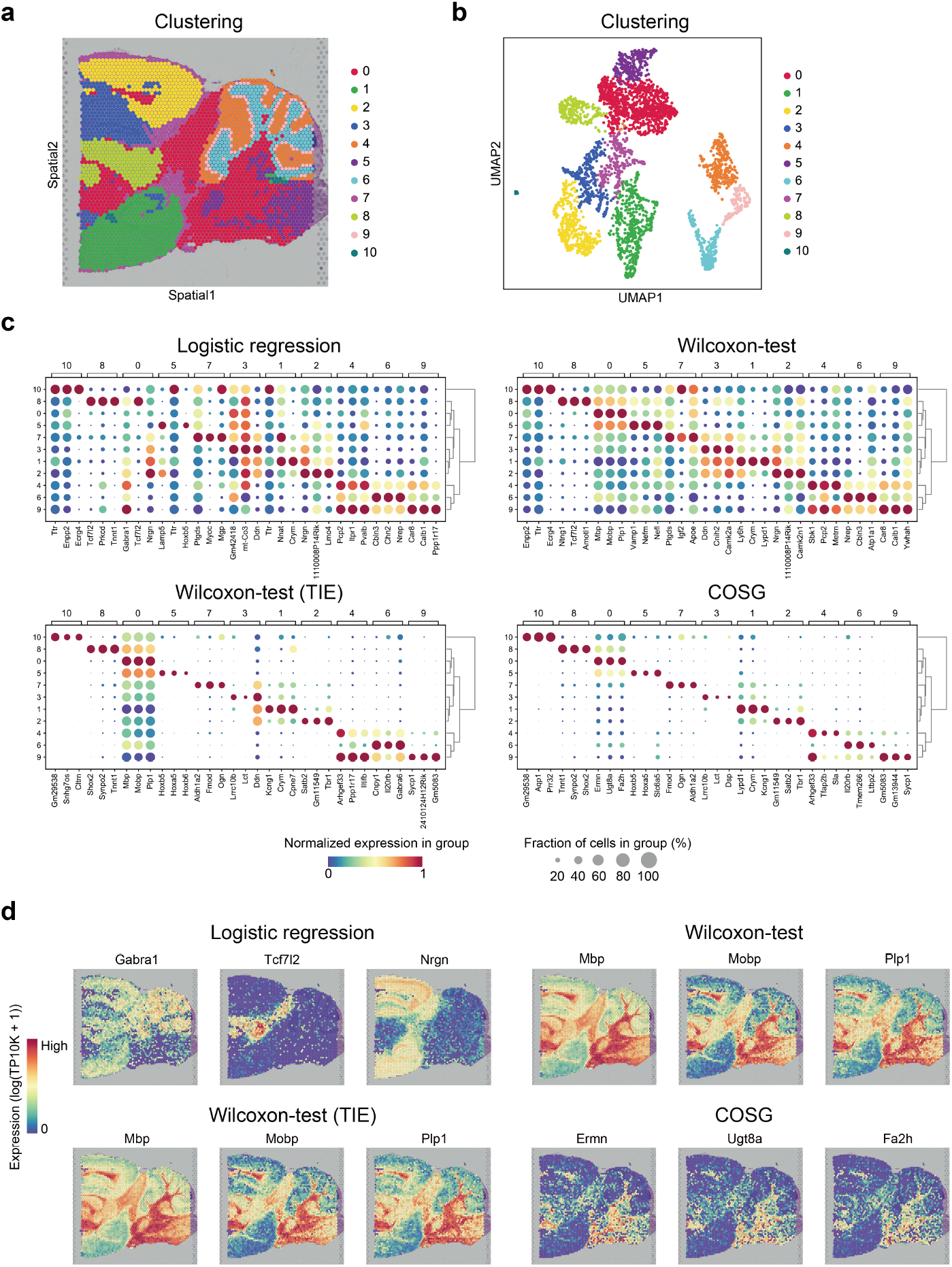
COSG performed well on spatial transcriptome data. **a**, Clustering results of the 3,355 signal spots detected in adult mouse brain sagittal posterior tissue. **b**, UMAP projection of the signal spots shown in (**a**). **c**, Expression dot plots of the top 3 marker genes identified by Logistic regression, Wilcoxon-test, Wilcoxon-test (TIE) and COSG for each cell cluster. **d**, Expression patterns of the top 3 marker genes identified by different methods for cells in Cluster 0.

We next analyzed the performance consistency of COSG across spatial transcriptomics platforms with a dataset generated by the Slide-seqV2 technology using mouse hippocampus^8^ (the Spatial-Slide-seqV2 dataset, Supplementary Table 3). The dataset contains 9,319 high-quality beads classified into 13 clusters (Supplementary Fig. 8a and 8b). Expression dot plots of the top 3 markers for each cell cluster demonstrated the superior performance of COSG as compared with other methods (Supplementary Fig. 8c). Expression pattern comparison of the top 3 marker genes for Cluster 5 showed that marker genes identified by COSG tended to have restricted expression in target cells, yet marker genes picked by other methods were broadly expressed (Supplementary Fig. 8d).

## Discussion

Marker gene identification safeguards the accuracy of cell type discrimination, therefore is a key step in single-cell sequencing data or spatial transcriptome data analysis. Here, we present COSG as a more accurate and faster method for marker gene identification from scRNA-seq, scATAC-seq and spatial transcriptome data. COSG should be applied to pre-clustered data to facilitate follow-up cell-type annotations, and the outputs of COSG can also be used to refine cell clustering results. COSG is implemented in both Python and R, and can be seamlessly used with popular toolkits, such as Scanpy^14^ and Seurat^15^.

The outstanding accuracy of COSG is achieved by assuming an ideal marker gene for each cell group and using cosine similarity to compare the expression patterns between the detected genes and the assumed ideal marker gene. Therefore, unlike other reported statistics-based marker gene identification methods, COSG is more robust to sequencing depth and capture efficiency of cells^13^, thus often generates more accurate results. Our experiments also showed that, due to the high frequency of missing values (zeros, or tied values), doing tie correction is necessary for Wilcoxon-test when it is applied to single-cell sequencing data.

COSG runs remarkably faster than other available methods, and it is capable of identifying marker genes from scRNA-seq data of over 1 million cells in less than 2 minutes. COSG is a universal method. It has achieved good performances in scATAC-seq and spatial transcriptome data, and also has the potential to be effectively applied to other types of single-cell omics data. The fast speed of COSG would be more beneficial when applying it to whole-genome scale single-cell sequencing data, as analysis of these types of data is usually time-consuming.

In short, COSG can serve as a general method for cell marker gene identification across different data modalities to facilitate single-cell data analysis and biomedical discoveries. Because the 10x Visium and Slide-seqV2 technologies are not at single-cell resolution yet, one spot or bead could contain several cells of multiple cell types, therefore the marker genes identified by COSG from spatial transcriptome data are not as discriminative as those from scRNA-seq data or scATAC-seq data. Enrichment analysis or aggregation of marker gene expressions may improve cluster annotations in spatial transcriptome data, which awaits future exploration.

## Methods

### Overview of COSG algorithm

COSG is designed to identify proper marker genes for predefined cell groups. The input data for COSG should first be normalized by other methods. After normalization, COSG generates the gene expression matrix *X* ∈ ℝ^*N*×*M*^, where *N* is the number of cells and *M* is the total number of detected genes. The *i*^th^ gene, ***g***_***i***_ ’s expression among all cells is the *i*^th^ column of *X*:

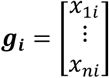

where *x*_*ji*_ is ***g***_***i***_ ’s expression value in the *j*^th^ cell, ***c***_***j***_, *j* ∈ {1, …, *N*}. Let *K* represents the number of cell groups predefined by manual annotation or unsupervised cell clustering. In order to identify marker genes for group *G*_*k*_, *k* ∈ {1, …, *K*}, we first set an ideal marker gene ***λ***_***k***_ for *G*_*k*_:

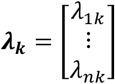

where *λ*_*jk*_ = 1 if ***c***_***j***_ ∈ *G*_*k*_ and *λ*_*jk*_ = 0 if ***c***_***j***_ ∉ *G*_*k*_.

We then calculate the cosine similarity between ***g***_***i***_ and ***λ***_***k***_ as *cos*(***g***_***i***_, ***λ***_***k***_):

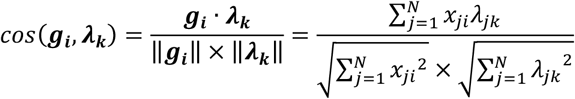

To evaluate whether ***g***_***i***_ is a good marker gene for group *G*_*k*_, we calculate COSG score as:

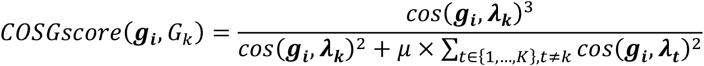

where *μ* ≥ 0 is the penalty factor for expression in non-target cell groups *G*_*t*_, *t* ∈ {1, …, *K*} and *t* ≠ *k*. The value of *μ* can be adjusted by users, and a larger *μ* means a bigger penalization for genes expressed in non-target cells. By default, *μ* = 1. The top marker genes for group *G*_*k*_ were selected by ranking COSG scores from the highest to the lowest.

### Methods compared with COSG

Besides COSG, 10 commonly used cell marker gene identification methods were evaluated in this study (Supplementary Table 2). Among them, Logistic regression was implemented by tl.rank_gene_groups (Scanpy v1.6.1) with ‘method’ set to ‘logreg’. Wilcoxon-test was implemented by tl.rank_gene_groups (Scanpy v1.6.1) with ‘method’ set to ‘wilcoxon’ and ‘tie_correct’ set to False. Wilcoxon-test (TIE) was implemented by tl.rank_gene_groups (Scanpy v1.6.1) with ‘method’ set to ‘wilcoxon’ and ‘tie_correct’ set to True. For t-test and t-test_overestim_var, we used tl.rank_gene_groups (Scanpy v1.6.1) with ‘method’ set to ‘t-test’ or ‘t-test_overestim_var’, respectively. For bimod, MAST, negbinom, poisson and roc, we used Seurat v3.2.3’s FindAllMarkers function with the parameter ‘test.use’ set to ‘bimod’, ‘MAST’, ‘negbinom’, ‘poisson’ or ‘roc’, respectively. All methods took normalized and log-transformed gene expression data as the input except for the negbinom and poisson methods, which took the raw count data as the input.

### Public data resources

#### scRNA-seq data

For the scRNA-seq dataset of mouse dentate gyrus cells^23^, the raw counts of unique molecular identifiers (UMIs) were downloaded from NCBI Gene Expression Omnibus (GEO) database (GSE104323). Data quality control was performed by filtering out genes detected in less than 3 cells and cells with any of the following features: 1) with fewer than 200 detected genes; 2) with more than 4,000 detected genes or more than 15,000 total UMIs; 3) with more than 20% UMIs derived from mitochondrial genomes. For the human kidney scRNA-seq data^24^, the preprocessed and normalized UMI counts were downloaded from the COVID-19 Cell Atlas (https://www.covid19cellatlas.org)^26^. **scATAC-seq data:** The raw scATAC-seq data and the processed genome track files of the mouse embryonic scATAC-seq dataset^4^ were downloaded from the NCBI GEO repository (GSE133244). The raw scATAC-seq data of the human bone marrow and peripheral blood mononuclear cells (BMMCs and PBMCs, respectively)^6^ was downloaded from https://github.com/GreenleafLab/MPAL-Single-Cell-2019 (File name: scATAC-Healthy-Hematopoiesis-191120.rds). **Spatial transcriptome data:** The adult mouse brain spatial transcriptome datasets (the sagittal posterior and coronal data) were downloaded from the 10x Genomics Visium spatial transcriptomics platform (https://support.10xgenomics.com/spatial-gene-expression/datasets/1.1.0/V1_Mouse_Brain_Sagittal_Posterior and https://support.10xgenomics.com/spatial-gene-expression/datasets/1.1.0/V1_Adult_Mouse_Brain). Genes detected in less than 3 spots were filtered out. The raw Slides-seqV2 mouse hippocampus spatial transcriptome data^8^ was downloaded from https://singlecell.broadinstitute.org/single_cell/study/SCP815/sensitive-spatial-genome-wide-expression-profiling-at-cellular-resolution (Puck_200115_08). Data quality control was performed by filtering out genes detected in less than 3 beads and beads met any of the following requirements: 1) with fewer than 200 detected genes or fewer than 1,000 total UMIs; 2) with more than 3,000 detected genes or more than 5,000 total UMIs; 3) with more than 20% UMIs derived from mitochondrial genomes.

### Generation of simulated datasets

The simulated datasets used in this study were generated by in-house built R scripts. Genes were simulated to follow negative binomial distribution using the rnbinom() function in R: *y* = rnbinom(n = *n*, mu = *μ*, size = 1), where n is the number of cells, mu is the mean expression value of each gene, and size is defined as the target number of successful trials. The following five types of gene expression patterns were simulated.

Type I expression represents the expression patterns of good marker genes, under which the simulated genes are specifically expressed in and restricted to target cells. For Type I genes, *μ* was set as 0.2 in target cells and 0.001 in non-target cells. Type II expression means the simulated genes are widely expressed, but with higher expression levels in target cells than in non-target cells. Genes with Type II expression pattern were simulated with *μ* set as 4 for 85% of the target cells and *μ* set as 2 for 85% of the non-target cells, and *μ* set as 2 or 4 for the remaining 15% of the target cells or the non-target cells, respectively. Type III expression means the simulated genes are not only expressed in the target group (*μ* = 0.4), but also expressed in limited numbers of non-target groups (3 non-target groups were created in each simulation, *μ* = 0.2 for each group). Type IV expression means the simulated genes have detectable but low expression in all cells, these genes were simulated with *μ* = 0.1. Type V expression means the simulated genes are highly expressed in all cells, these genes were simulated with *μ* = 2.

Using the above procedure, we generated 30 simulated datasets (each contains 20 cell groups). The number of cells contained in the simulated datasets ranged from 1,000 to 10,000, with the number of cells per dataset increased 1,000 per step. At each total cell number, three datasets were generated with different population distributions among cell groups. For all datasets, the minimum number of cells for a cell group was set as 5. A total of 20 genes with Type I expression pattern were generated as the real marker genes for each cell group. In addition, 20 genes with Type II expression and 20 genes with Type III expression were generated for each cell group to serve as the confounding factors for the real marker genes. For all cell groups in each dataset, the numbers of genes with Type IV expression and genes with Type V expression were both set as 500.

### Generation of large-scale experimental benchmark datasets

To generate large-scale benchmark datasets, we subsampled the Drop-seq scRNA-seq dataset of *Tabula Muris Senis*^27^ (TMS, 245,389 cells of 123 annotated cell types) to generate 14 experimental benchmark datasets (each contained 31 cell types) with sizes ranging from 1,000 to 150,000 cells (Supplementary Table 4). The raw UMI count data was downloaded from https://figshare.com/articles/dataset/tms_gene_data_rv1/12827615?file=24351014. Cell types with too few cells (less than 2,000 cells) or too many cells (more than 30,000 cells) were filtered out to avoid sampling bias. The remaining 156,630 cells from 31 cell types were subsampled using the pp.subsample function (Scanpy v1.6.1). Cell replacement was not allowed during the subsampling process.

To generate benchmark datasets from the Mouse Organogenesis Cell Atlas^28^, the filtered high-quality scRNA-seq UMI count data (File name: gene_count_cleaned.RDS) was downloaded from the Mouse Organogenesis Cell Atlas website (https://oncoscape.v3.sttrcancer.org/atlas.gs.washington.edu.mouse.rna/downloads). The downloaded data has 1,331,984 cells with annotations. Genes detected in less than 3 cells were filtered out. Benchmark datasets with 50,000, 100,000, 500,000, 1,000,000 and 1,331,984 cells, respectively, were generated using the same method mentioned above. Each dataset contains exactly 37 cell types.

### Data normalization

Except for the human kidney scRNA-seq dataset which used the normalized UMI counts provided by the dataset owners, all other datasets were normalized using the below methods. For the scRNA-seq and spatial transcriptome data, normalization was performed by firstly dividing the raw counts of each gene within each cell/spot/bead by the total number of raw counts within that cell/spot/bead, and then multiplying by 10,000, using the pp.normalize_total function in Scanpy v1.6.1. The normalized counts were then log-transformed via the pp.log1p function in Scanpy v1.6.1. For the scATAC-seq data, the raw scATAC-seq data was normalized by the term frequency-inverse document frequency (TF-IDF) algorithm implemented by the RunTFIDF function of Signac v1.1.0^29^.

### Dimensionality reduction

We used Principal Component Analysis (PCA) to embed the detected cells/spots/beads of scRNA-seq or spatial transcriptome data into a low-dimensional space. Before PCA, we selected 3000 highly-variable genes by the pp.highly_variable_genes function (Scanpy v1.6.1) and used them as the input data for PCA. We first used the StandardScaler function of scikit-learn v0.24.0^30^ to scale the highly-variable genes to unit variance to diminish the effects of gene abundance difference, then applied the TruncatedSVD function of scikit-learn v0.24.0 to obtain PCA embedding. The default component number of PCA was set as 50. The PCA embedding results were used for downstream UMAP visualization and Leiden clustering.

### Data visualization

The 2-dimensional distributions of cells/spots/beads within each dataset were visualized by Uniform Manifold Approximation and Projection (UMAP)^31^ plots using the top 50 principal components (PCs) of the scRNA-seq datasets and the top 30 PCs of the spatial transcriptome datasets. UMAP was implemented via the pp.neighbors function (parameters: n_neighbors=15, knn=True, use_rep =‘X_pca’ and method=‘umap’) followed by the tl.umap function (with default parameters) of Scanpy (v1.6.1). For the human kidney scRNA-seq dataset^24^ and the scATAC-seq datasets^4,6^, the 2-dimensional coordinates of cells were adopted from the original publications and used for UMAP plot construction.

### Cell type annotation

For the scRNA-seq and scATAC-seq datasets, the identities of cells were characterized according to the original publications^4,6,23,24,27,28^. For the spatial transcriptome datasets, unsupervised graph-based Leiden clustering algorithm^32^ implemented by the pp.neighbors (parameters: n_neighbors=15, knn=True, use_rep=‘X_pca’ and method=‘umap’) and tl.leiden functions (Scanpy v1.6.1) were used to cluster spots/beads into different groups according to their gene expression similarities. The resolution parameter for tl.leiden (Scanpy v1.6.1) was set as 0.3 for the Spatial-brain_sagitta dataset, set as 0.25 for the Spatial_brain_coronal dataset and set as 0.5 for the Slide-seqV2 spatial transcriptome dataset.

### Running time evaluation

The running time for each tested marker gene identification method was measured by the time module in Python. All methods were run on a 2.00GHz Intel Xeon E7-4830v4 central processing unit (CPU) with 512GB of RAM. Except for the MAST method, which by default uses multiple CPU cores, other methods were restricted to use one CPU core.

## Code availability

COSG is available at https://github.com/genecell/COSG (Python) and https://github.com/genecell/COSGR (R).

## Acknowledgements

This work was supported by the National Key Research and Development Program of China (2019YFC1708903), Natural Science Foundation of China (81790622 and 91940304), CAS Strategic Priority Research Program (XDA16020801), Beijing Natural Science Foundation of China (Z200020) to X.-J. W.

## Author contributions

XW and XP supervised the study. MD developed the algorithm, built the computational tools, and performed the analysis. XW and MD wrote the manuscript. All authors read and approved the final manuscript.

---

**Supplementary Fig. 1.**
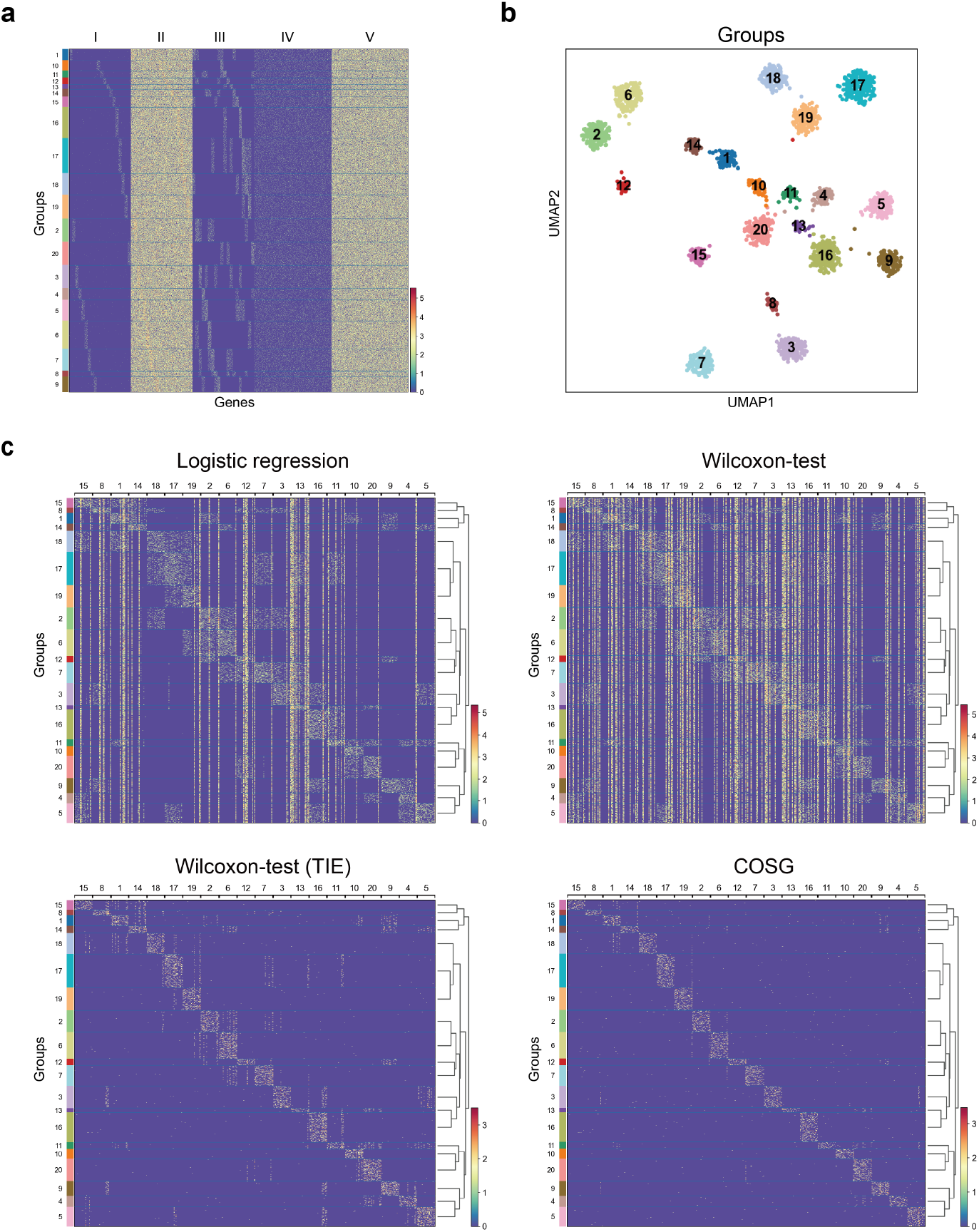
Gene expression patterns in the simulated scRNA-seq dataset. **a**, Gene expression heatmap shows the five simulated patterns of gene expression among 2,000 cells. **b**, UMAP projection of the simulated scRNA-seq data with 2,000 cells. Colors represent different cell groups. **c**, Gene expression heatmap shows the expression patterns of the top 20 marker genes for each cell group identified by Logistic regression, Wilcoxon-test, Wilcoxon-test (TIE) and COSG.

**Supplementary Fig. 2.**
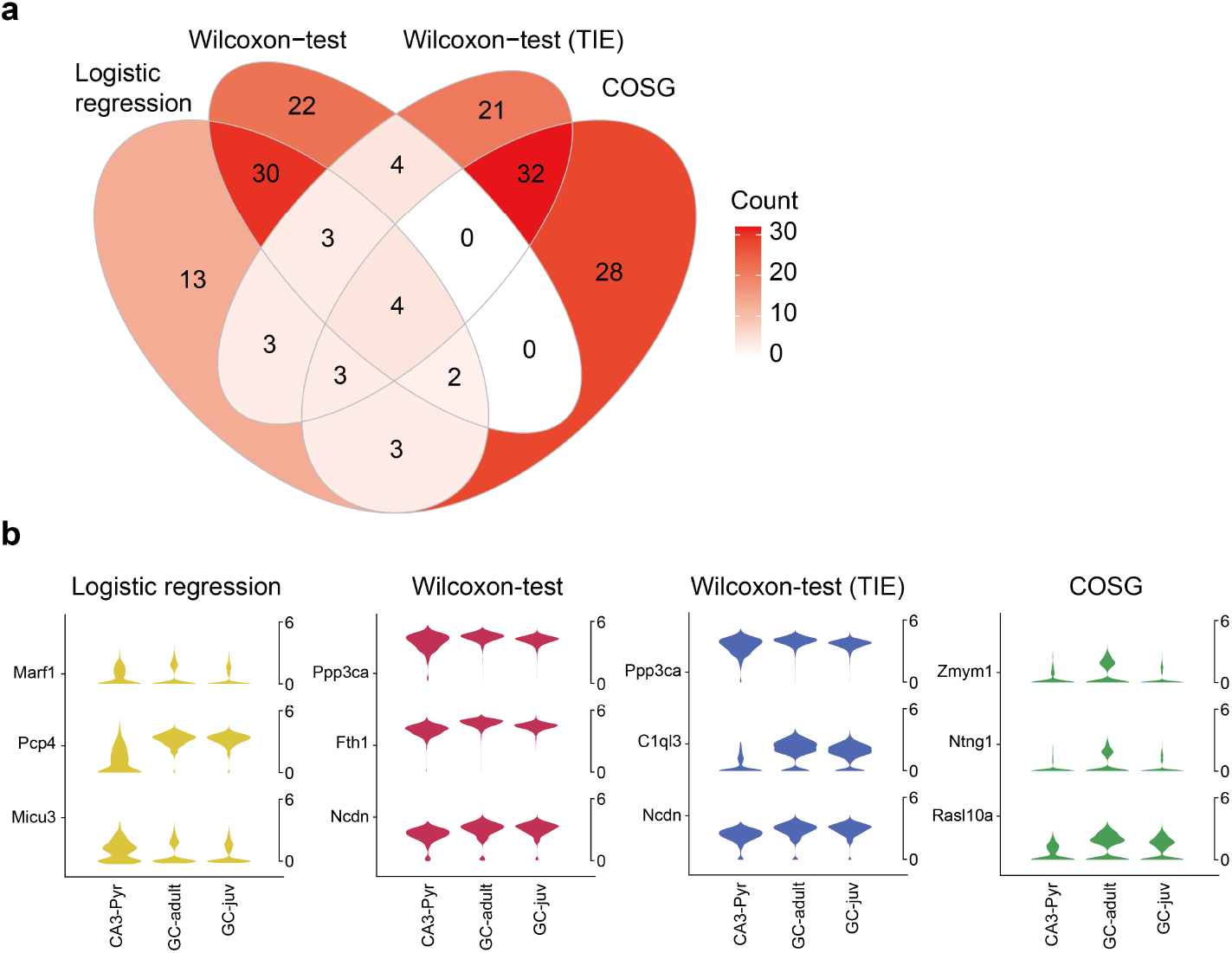
Marker genes identified by COSG for the RNA-Hochgerner dataset are more indicative than those identified by other methods. **a**, Venn diagram of the joint set of the top 3 marker genes identified by Logistic regression, Wilcoxon-test, Wilcoxon-test (TIE) and COSG for each cell type in the RNA-Hochgerner dataset. **b**, Violin plots representing the normalized expression values of the top 3 marker genes identified by each method for GC-adult cells. GC, granule cell; CA3, hippocampus CA3 pyramidal layer; Pyr, pyramidal cell; juv, juvenile.

**Supplementary Fig. 3.**
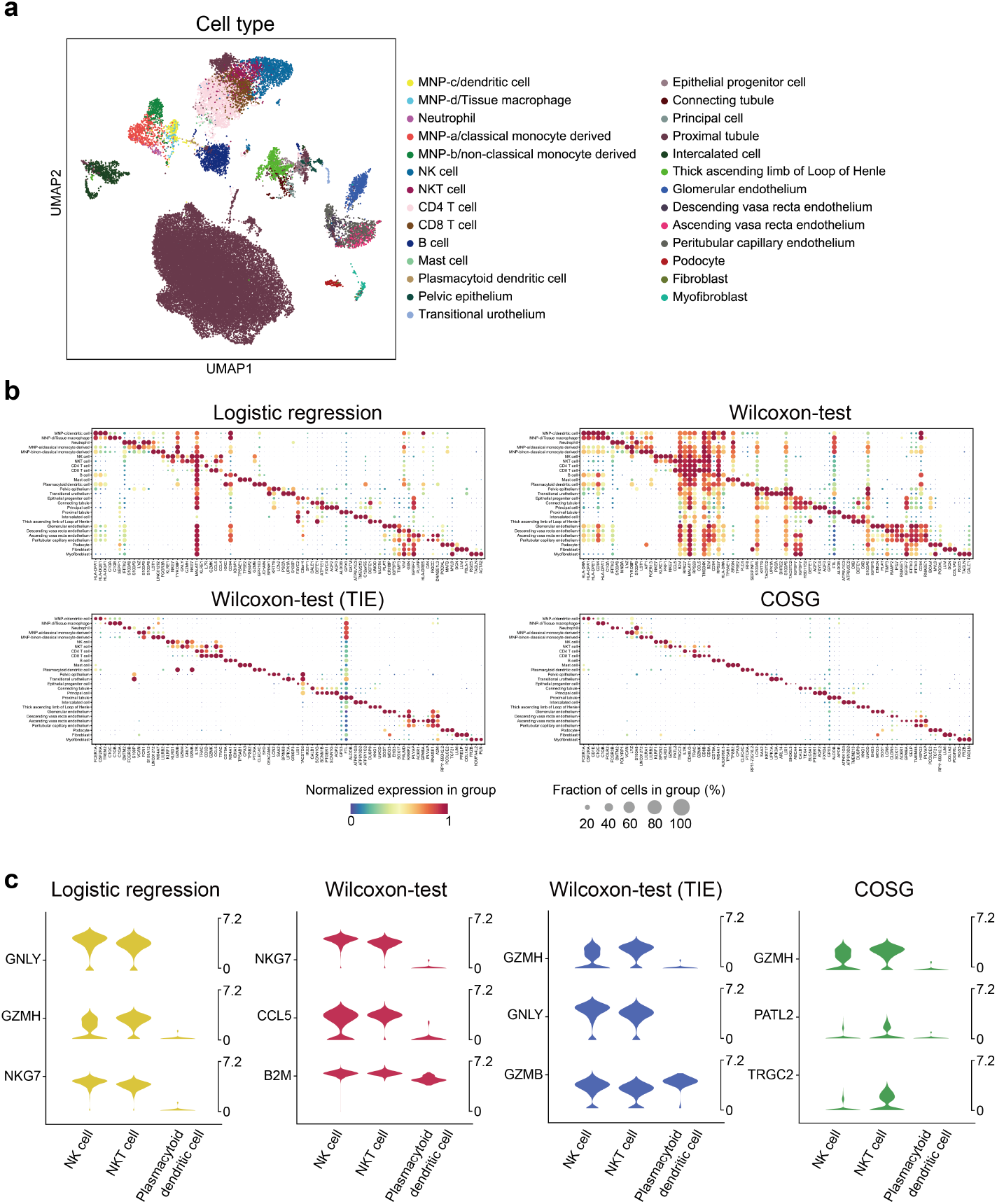
COSG outperforms other methods on the RNA-Stewart dataset. **a**, UMAP projection of the scRNA-seq data of 40,268 human adult kidney cells (the RNA-Stewart dataset). **b**, Expression dot plots of the top 3 marker genes identified by Logistic regression, Wilcoxon-test, Wilcoxon-test (TIE) and COSG for each cell type. **c**, Violin plots representing the normalized expression values of the top 3 marker genes identified by each method for NKT cells. NK cell and plasmacytoid dendritic cell are included for comparison.

**Supplementary Fig. 4.**
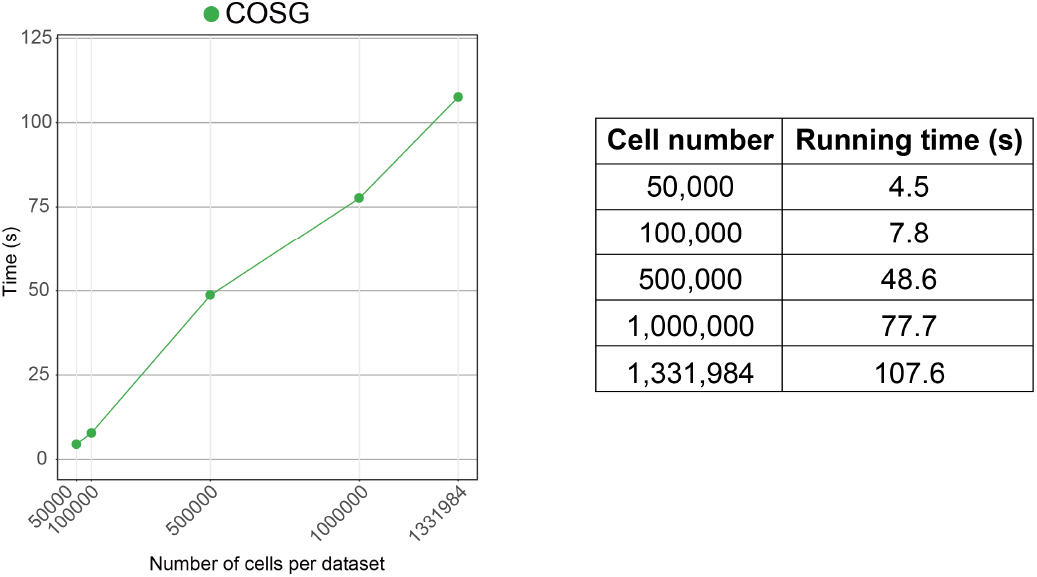
Running time of COSG on the million-scale Mouse Organogenesis Cell Atlas dataset. The Mouse Organogenesis Cell Atlas dataset has 1,331,984 annotated cells. Benchmark datasets with 50,000, 100,000, 500,000, 1,000,000 and 1,331,984 cells were generated and used to test the efficiency of COSG.

**Supplementary Fig. 5.**
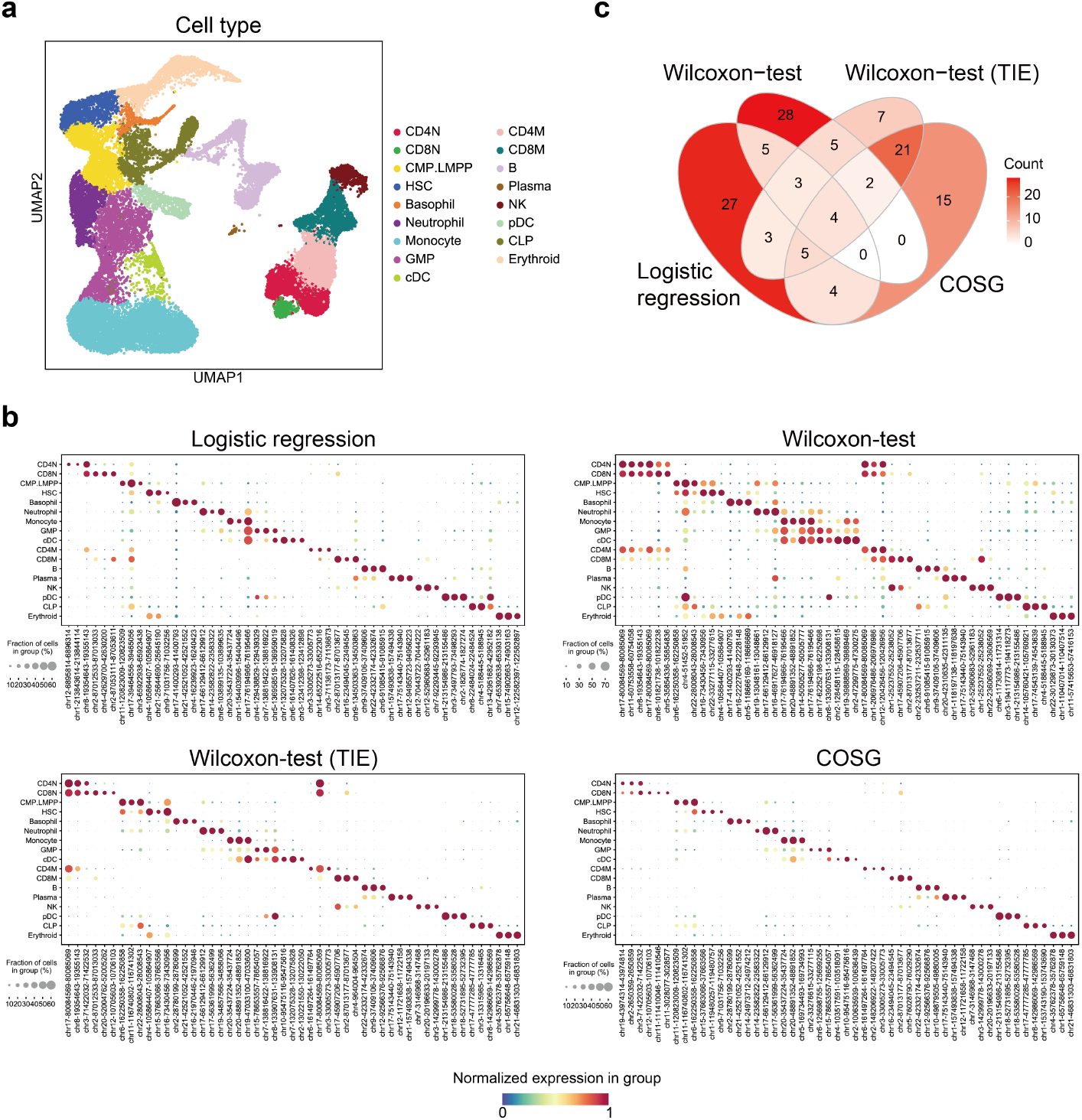
Marker regions identified by COSG from the ATAC-Granja_broad dataset have better cell type specificity. **a**, UMAP projection of the scATAC-seq data of 33,819 human bone marrow cells and PBMCs (17 cell types). **b**, Expression dot plots of the top 3 marker regions identified by Logistic regression, Wilcoxon-test, Wilcoxon-test (TIE) and COSG for each cell type. **c**, Venn diagram of the joint set of the top 3 marker regions for each cell type identified by different methods.

**Supplementary Fig. 6.**
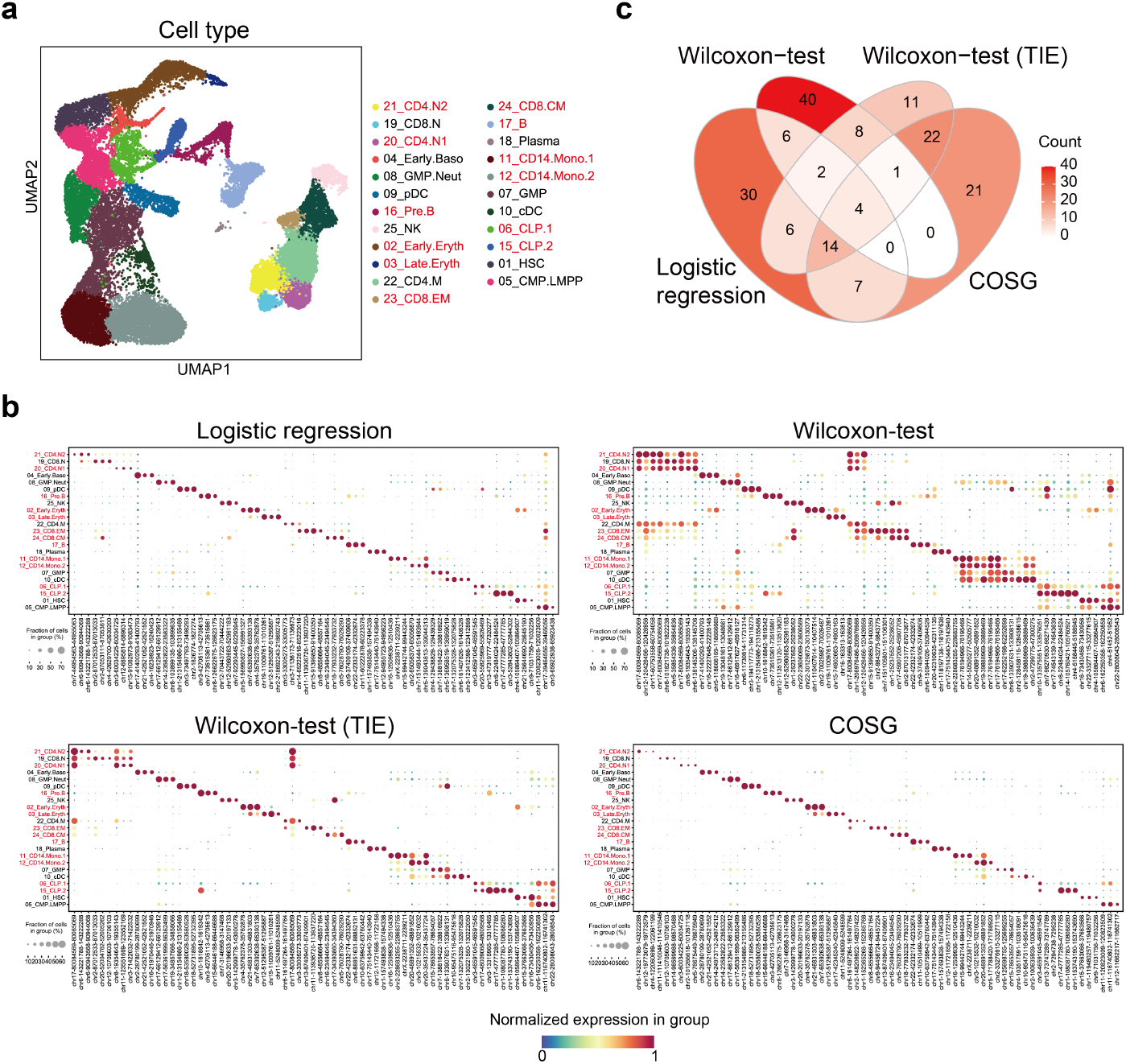
Marker regions identified by COSG from the ATAC-Granja_fine dataset are more indicative than those identified by other methods. **a**, UMAP projection of the scATAC-seq data of 33,819 human bone marrow cells and PBMCs (23 cell types). **b**, Expression dot plots of the top 3 marker regions identified by Logistic regression, Wilcoxon-test, Wilcoxon-test (TIE) and COSG for each cell type. **c**, Venn diagram of the joint set of the top 3 marker regions for each cell type identified by different methods. In (**a**) and (**b**), names of extra cell types not included in Supplementary Fig. 5 are shown in red.

**Supplementary Fig. 7.**
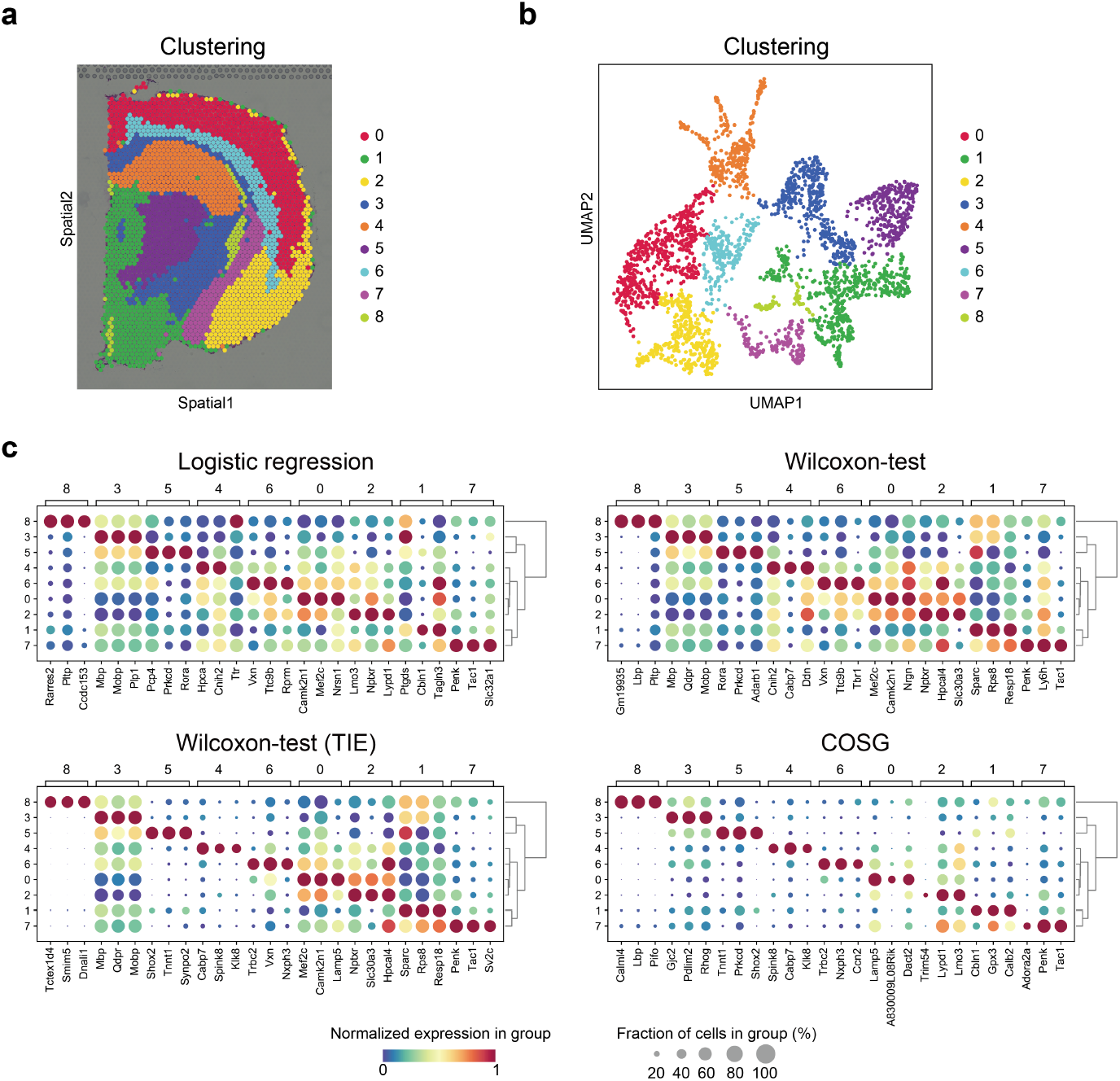
COSG performed well on the Spatial-brain_coronal dataset. **a**, Clustering results of the 2,702 signal spots detected in adult mouse brain coronal tissue. **b**, UMAP projection of signal spots shown in (**a**). **c**, Expression dot plots of the top 3 marker genes for each cluster identified by different methods.

**Supplementary Fig. 8.**
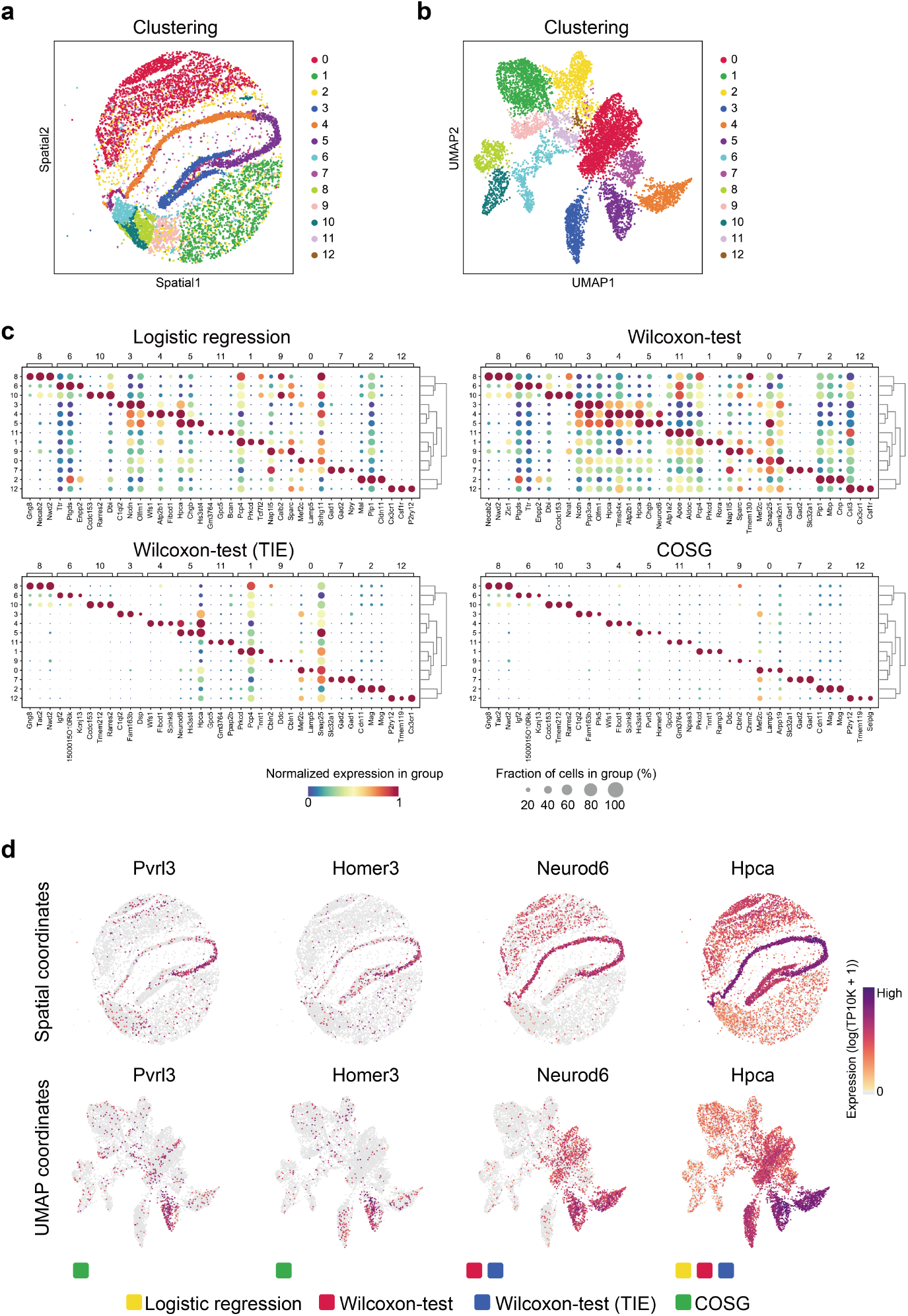
COSG outperformed other methods on the Spatial-Slide-seqV2 dataset. **a**, Clustering results of the 9,319 beads obtained from a section of mouse hippocampus. **b**, UMAP projection of the detected beads in (**a**). **c**, Expression dot plots of the top 3 marker genes for each cluster identified by different methods. **d**, Gene expression patterns of the top marker genes for cells in Cluster 5 identified by different methods.

**Supplementary Table 1.** List of 30 simulated datasets generated by the simulation procedure and used for the accuracy benchmark testing in this study. Each dataset contains 20 cell groups.

| No. of cells | No. of replicates |
| --- | --- |
| 1,000 | 3* |
| 2,000 | 3 |
| 3,000 | 3 |
| 4,000 | 3 |
| 5,000 | 3 |
| 6,000 | 3 |
| 7,000 | 3 |
| 8,000 | 3 |
| 9,000 | 3 |
| 10,000 | 3 |

\*Note: the 3 replicates each has different ratios of cell groups.

**Supplementary Table 2.** List of 11 marker gene identification methods tested in this study.

| Name of method | Algorithm | Implementation (version) |
| --- | --- | --- |
| 1 COSG | Cosine similarity-based method | Python package COSG (1.0.0) |
| 2 Logistic regression | Logistic regression | Python package Scanpy (1.6.1) |
| 3 Wilcoxon-test | Wilcoxon Rank Sum test (without tie correction) | Python package Scanpy (1.6.1) |
| 4 Wilcoxon-test (TIE) | Wilcoxon Rank Sum test (with tie correction) | Python package Scanpy (1.6.1) |
| 5 t-test | Student's t-test | Python package Scanpy (1.6.1) |
| 6 t-test_overestim_var | Student's t-test that overestimates variance of each group | Python package Scanpy (1.6.1) |
| 7 MAST | Hurdle model | R packages MAST (1.12.0) and Seurat (3.2.3) |
| 8 bimod | Likelihood-ratio test | R package Seurat (3.2.3) |
| 9 negbinom | Negative binomial generalized linear model | R package Seurat (3.2.3) |
| 10 poisson | Poisson generalized linear model | R package Seurat (3.2.3) |
| 11 roc | ROC classifier | R package Seurat (3.2.3) |

**Supplementary Table 3.** List of single-cell sequencing datasets used in this study.

| Dataset | Description | Methods | Dataset size | No. of features | No. of groups | Reference |
| --- | --- | --- | --- | --- | --- | --- |
| RNA-Hochgerner | Dentate gyrus cells in perinatal, juvenile, and adult mice | Droplet scRNA-seq (10x Genomics) | 23,025 cells | 19,444 genes | 24 | Hochgerner et al., *Nat. Neurosci.*, 2018 |
| RNA-Stewart | The spatiotemporal immune topology of human adult kidney | Droplet scRNA-seq (10x Genomics) | 40,268 cells | 33,694 genes | 27 | Stewart et al., *Science*, 2019 |
| RNA-TMS_drop | Multiple mouse tissues and organs | Microfluidic droplet | 150,000 cells | 20,138 genes | 31 | Tabula Muris Consortium, *Nature*, 2019 |
| ATAC-Pijuan-Sala | Single nuclei from mouse embryos at 8.25 days post-fertilization | Single-nucleus ATAC-seq | 19,453 cells | 301,316 genomic regions | 18 | Pijuan-Sala et al., *Nat. Cell Biol.*, 2020 |
| ATAC-Granja_broad | Human healthy bone marrow and PBMCs | Droplet scATAC-seq (10x Genomics) | 33,819 cells | 451,999 genomic regions | 17 | Granja et al., *Nat. Biotechnol.*, 2019 |
| ATAC-Granja_fine* | Human healthy bone marrow and PBMCs | Droplet scATAC-seq (10x Genomics) | 33,819 cells | 451,999 genomic regions | 23 | Granja et al., *Nat. Biotechnol.*, 2019 |
| Spatial-brain_sagitta | Mouse brain (sagittal posterior) | 10x Visium | 3,355 spots | 19,147 genes | 11 | 10x Genomics |
| Spatial-brain_coronal | Mouse brain (coronal) | 10x Visium | 2,702 spots | 19,652 genes | 9 | 10x Genomics |
| Spatial-Slide-seqV2 | Mouse hippocampus | Slide-seqV2 | 9,319 beads | 20,424 genes | 13 | Stickels et al., *Nat. Biotechnol.*, 2020 |

654 Note: \*The ATAC-Granja\_broad dataset and the ATAC-Granja\_fine dataset have the same gene expression profiles, but different cell group numbers due to different annotation depths.

**Supplementary Table 4.** List of datasets subsampled from the *Tabula Muris Senis* (Drop-seq) dataset and used for the running time benchmark testing in this study.

| Dataset | No. of cells | No. of cell types | Tested methods |
| --- | --- | --- | --- |
| D1 | 1,000 | 31 | A + B |
| D2 | 2,000 | 31 | A + B |
| D3 | 3,000 | 31 | A + B |
| D4 | 4,000 | 31 | A + B |
| D5 | 5,000 | 31 | A + B |
| D6 | 6,000 | 31 | A + B |
| D7 | 7,000 | 31 | A + B |
| D8 | 8,000 | 31 | A + B |
| D9 | 9,000 | 31 | A + B |
| D10 | 10,000 | 31 | A + B |
| D11 | 20,000 | 31 | B* |
| D12 | 50,000 | 31 | B* |
| D13 | 100,000 | 31 | B* |
| D14 | 150,000 | 31 | B* |

\*Basing on the results of D1 to D10, only the fastest six methods were run on larger datasets containing more than 10,000 cells. A and B denote different groups of methods. A: bimod, MAST, negbinom, poisson and roc. B: t-test, t-test\_overestim\_var, Wilcoxon-test, Logistic regression, Wilcoxon-test (TIE) and COSG.

**Supplementary Table 5.**
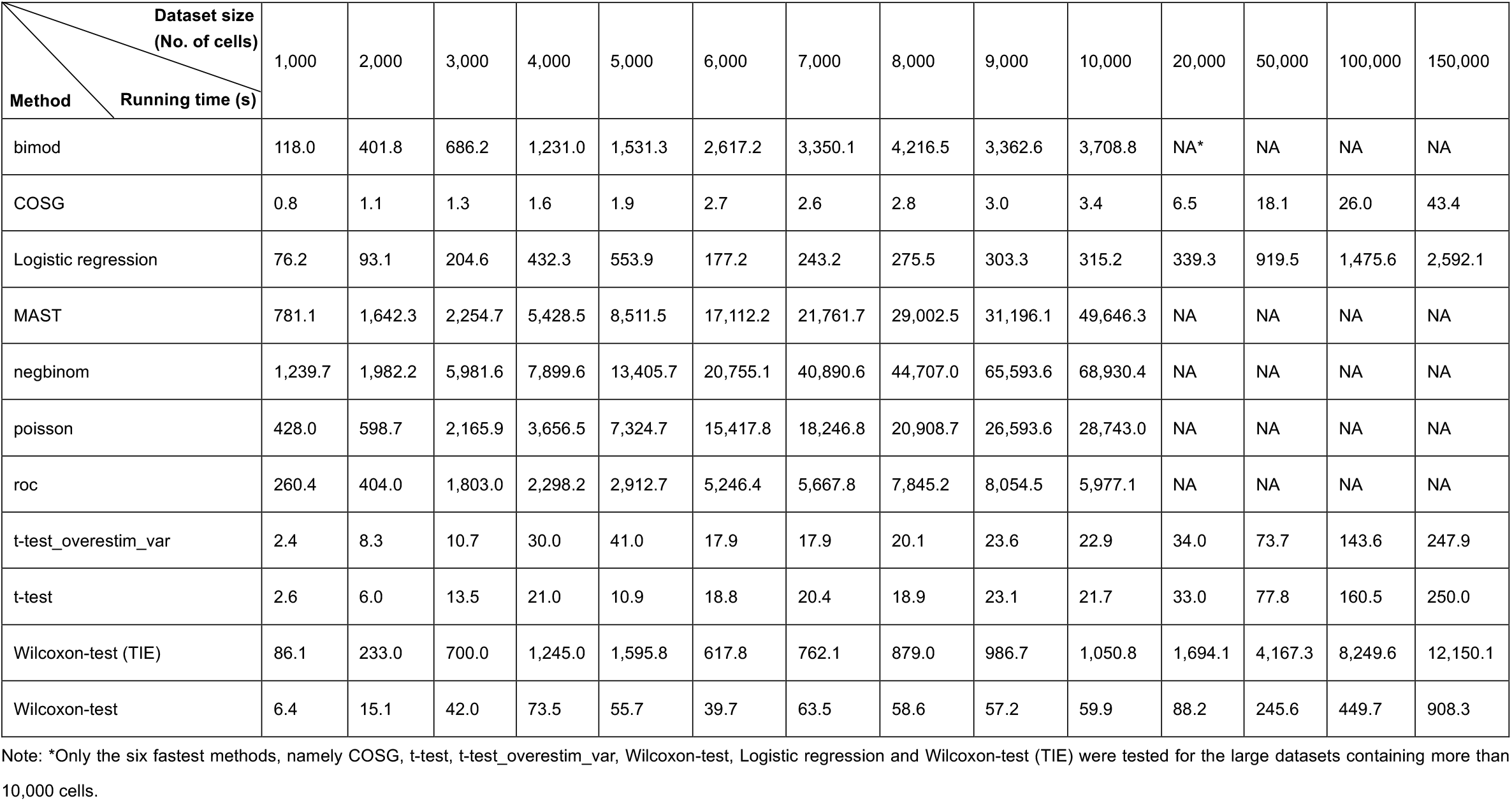
Running time (seconds) of 11 methods on datasets with cell numbers ranging from 1,000 to 150,000. These experimental benchmark datasets were subsampled from the Drop-seq scRNA-seq dataset of *Tabula Muris Senis*.

**Supplementary Table 6.**
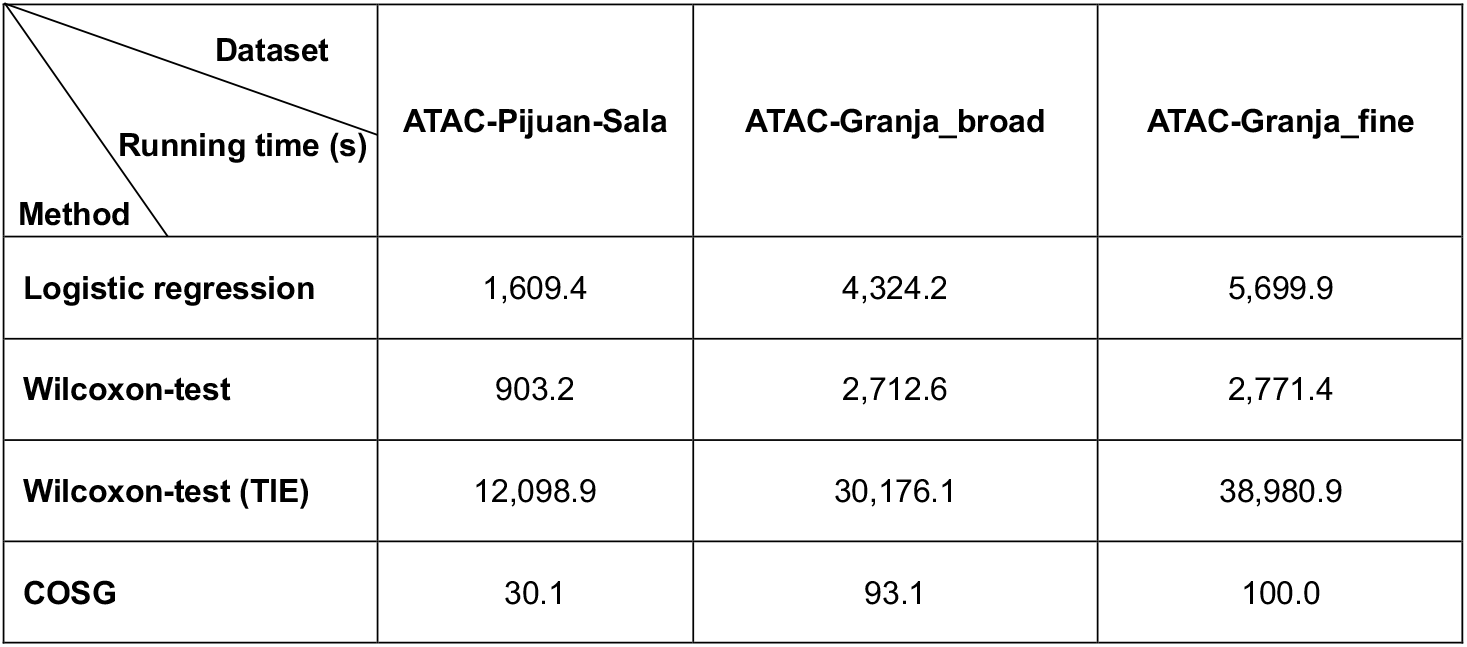
Running time (seconds) of Logistic regression, Wilcoxon-test, Wilcoxon-test (TIE) and COSG on the three scATAC-seq datasets.

| Dataset<br>Running time (s)<br>Method | ATAC-Pijuan-Sala | ATAC-Granja_broad | ATAC-Granja_fine |
| --- | --- | --- | --- |
| Logistic regression | 1,609.4 | 4,324.2 | 5,699.9 |
| Wilcoxon-test | 903.2 | 2,712.6 | 2,771.4 |
| Wilcoxon-test (TIE) | 12,098.9 | 30,176.1 | 38,980.9 |
| COSG | 30.1 | 93.1 | 100.0 |

